# Spectrally Resolved Localization Microscopy with Ultrahigh-Throughput

**DOI:** 10.1101/2024.01.17.576082

**Authors:** James Ethan Batey, Geun Wan Kim, Meek Yang, Darby Claire Heffer, Elric Dion Pott, Hannah Giang, Bin Dong

## Abstract

Single-molecule localization microscopy (SMLM) has become a strong technique in the toolbox of chemists, biologists, physicists, and engineers in recent years for its unique ability to resolve characteristic features quickly and accurately in complex environments at the nanoscopic level. Multicolor super-resolution imaging has seen the greatest advancement among SMLM techniques, drastically improving the differentiation ability of nanostructures beyond the diffraction limit and increasing the resolution with which previously unresolvable structures are studied. However, current multicolor SMLM methodologies present low spatial resolution and throughput and require complex optical systems. Here, we overcome these drawbacks by developing an ultrahigh-throughput SMLM methodology that allows for ultrahigh throughput multicolor imaging at the nanoscopic level using a color glass filter. Our methodology can readily distinguish fluorophores of close spectral emission and achieves sub-10 nm localization and sub-5nm spectral precisions.

## Introduction

Super-resolved fluorescence microscopy has gained widespread attention across multiple research areas^1-3^ due to its ability to resolve structures with spatial resolution beyond the diffraction limit. Especially single-molecule localization microscopy (SMLM)^4^ has been at the forefront of super-resolution methodologies in the last decades. In this technique, stochastic single fluorophore emission creates well-separated individual point spread functions (PSFs). Modern SMLM methods such as stochastic optical reconstruction microscopy (STORM),^5^ photoactivated localization microscopy (PALM),^6^ and point accumulation for imaging in nanoscale topography (PAINT)^7^ have advanced in their abilities to precisely localize single fluorophore PSFs and resolve structural features with low computational requirements, simple instrumental and experimental setups. The level of spatial resolution provided by SMLM techniques reveals previously unresolvable kinetic, dynamic, and structural information of sophisticated biological,^1, 4^ chemical,^2^ and material samples^8, 9^ that exist beyond the diffraction limit thus enabling application at the subcellular and molecular level.

Spectro-microscopic systems have been developed for simultaneously acquiring high-resolution images and spectral features at the single-molecule level.^10-12^ These techniques make use of dichroic filter,^13-15^ PSF engineering,^16, 17^ and spectral dispersion.^10-12, 18-21^ First, ratiometric detection employed one or more appropriate dichroic mirrors (DCMs) to distinguish fluorophores based on their respective emission intensity in two channels. In this approach, localization precision suffers when more than two colors are detected, as significant signal loss arises when adding multiple DCMs. Furthermore, The only spectral information obtainable by using DCMs is whether a fluorophore has a wavelength greater or lesser than the DCM’s respective cutoff band.^6^ Second, PSF engineering can distinguish fluorophores with different colors by inserting a spatial light modulator (SLM) or a phase mask (PM) in the Fourier plane of the emission path. The resulting PSFs shape depends on the emission wavelength. However, this method has low spectral resolution. Fluorophores that have small differences in emission wavelengths (i.e., in the order of tens of nanometers) cannot be differentiated. This method has challenges at high density of fluorescent emitters where severe interference on the PSFs shapes presents. Third, in spectral dispersion method, dispersive optical elements (e.g., diffraction grating or prism) spatially separate and project the spatial and spectral components of the emission signal on different parts of the detector.^10^ This method allows the full emission spectra from single molecules to be obtained. However, its spatial resolution is limited because most photons are used for spectral analysis while only a small fraction of photons for obtaining fluorophores’ spatial positions. It also requires low molecular density to eliminate spectral interference for fluorophores that are in close spatial proximity. These three implementations present significant challenges for studying fluorogenic reactions^22^, subcellular mechanisms^23-26^, and other applications that benefit from high-throughput SMLM techniques with high spatial and spectral resolution.

Here, we developed an imaging methodology capable of overcoming the challenges associated with current multicolor SMLM techniques. A two-channel imaging module employing a 50:50 beam splitter was custom-built and integrated into a total internal reflection fluorescence microscope. The fluorescence emission in one channel was modified by a color glass filter (CGF). A correlation between obtained photon intensities in two channels with the wavelength was established, which enables the determination of fluorophores’ emission spectral mean. In this approach, the throughput of single molecule localization analysis is the same as that in single color SMLM. Furthermore, the photons in both channels are combined for determining fluorophores’ positions with increased spatial resolution. We further demonstrate that our method can distinguish fluorophores with heavily overlapping emission spectra. With the implementation of a simple optical component, we show that spectrally resolved SMLM can be achieved with ultrahigh-throughput, sub-10 nm spatial and sub-5 nm spectral precisions.

## Results and Discussions

### Ultrahigh throughput single molecule spectroscopy and microscopy imaging

To spectrally resolve and spatially localize single molecules with the highest attainable throughput, the signal used for spectral measurement and localization needs to be spatially far-off from each other. To accomplish this, we designed and built a custom dual-channel detection box and equipped it with a high transmission efficiency nonpolarizing 50:50 beam splitter (Figure 1a). Two identical light paths were created as half the fluorophore emission was reflected and the other half was transmitted through the beam splitter. A color glass filter (CGF, Figure S1) was then placed into the transmitted light path. It modifies the signal in a way that lowers its total intensity analogous with how closely the fluorophore’s emission spectrum matches the CGF’s transmission window. The reflected signal out of the beam splitter remains unmodified. Both the modified and unmodified signals were focused side-by-side on the same camera chip. Notably, this allows for ultrahigh throughput to be achieved on a single camera chip. Alternatively, multiple cameras can be used in this implementation to expand the field of view.

**Figure 1.**
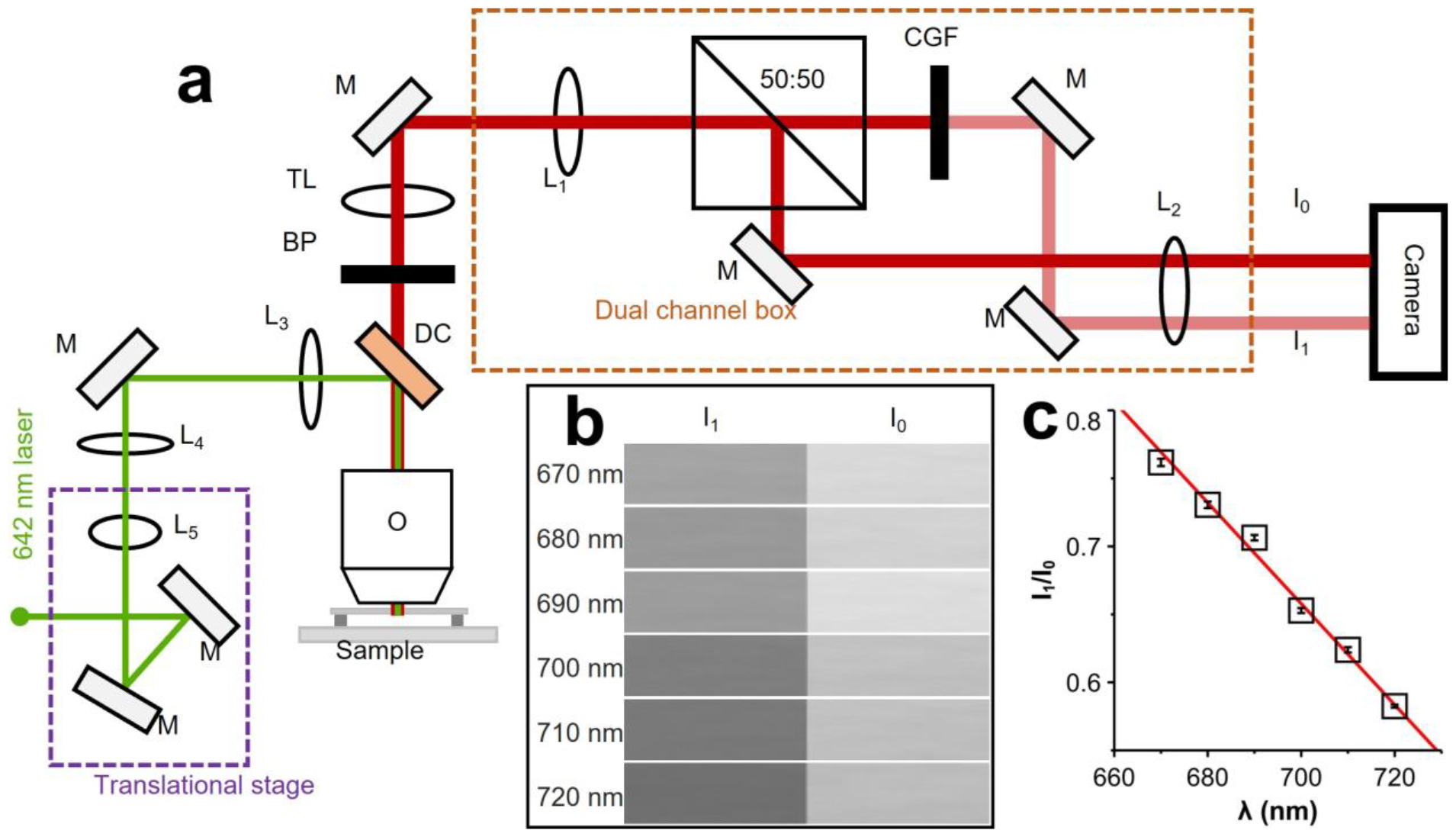
Single molecule spectroscopy and microscopy imaging. (a) schematic view of the imaging setup. (b) Bright field images of light source at different wavelengths. The light source was filtered with short band pass filters before projecting to the sample. (c) Wavelength dependence of ratio of intensity (I_1_/I_0_) in CGF and reference channel.

The side-by-side channels were propagated onto the camera so that the left channel contains the CGF modified signal and the right channel contains the reference signal (Figure 1b). The modified channel’s intensity is lower than that of the reference channel. The amount by which the modified signal intensity is reduced by the CGF depends on the emission profile of the fluorophore (Figure S1). The ratio of modified signal intensity (I_1_) to unmodified signal intensity (I_0_) is a unique value that changes exclusively with wavelength. We identified a relationship between I_1_/I_0_ ratio and wavelength directly using the light source equipped in the microscope. The light source was first filtered by short bandpass filters (670 – 720 nm) and then projected to the camera after passing the dual-channel detection box. Figure 1b shows the dual channel bright-filed images of the light source at different wavelengths. Both intensities (I_1_ and I_0_) were obtained from corresponding images as shown in Figure 1b. At each wavelength, 500 frames of images were taken, and their intensities were averaged to find the mean I_1_/I_0_ ratio. The correlation between I_1_/I_0_ ratio and wavelength is displayed in Figure 1c and shows a highly linear relationship. Using this obtained calibration curve, the spectral mean of a fluorophore can be determined from its determined I_1_/I_0_ ratio.

We first tested this method with Alexa Fluor 647 (AF647), one of the most used dyes for STORM imaging. We fixed and labeled the microtubule filaments of A549 cells with AF647. We applied the direct STORM approach^27^ by photoswitching the majority of the dye molecules into nonfluorescent dark state, leaving a random subset and spatially resolved dye molecules fluorescing (on state) in each imaging frame (Figure 2a). The single molecules in both channels were first identified and localized independently. Then correlation analysis (Figure S2) was applied to identify the same single molecules in both channels and their photon intensities were used for calculating the I_1_/I_0_ ratio. Applying the calibration curve described above, we obtained the spectral mean (λ_mean_). Figure 2b and c shows the histogram distribution of I_1_/I_0_ ratio and λ_mean_ from millions of AF647 dye molecules, respectively. Fitting the distribution data gives λ_mean_ of 689.2 ± 5.3 nm. We repeated the experiments with another photoswitchable dye, cyanine 5.5 (Cy 5.5), and obtained λ_mean_ of 707.9 ± 5.5 nm (Figure S3). Combining the obtained single molecule spatial coordinates and the spectral mean, spectrally resolved super-resolution fluorescence images (SR-SMLM) can then be reconstructed (Figure 2d and Figure S3). Though the statistical analysis shows narrow distribution of λ_mean_, heterogeneity of emission spectra is clear observed from the SR-SRFM image. We further evaluate the spatial resolution of our imaging system through cluster analysis (Figure S4). Similar localization precisions (σ) of ∼10 nm, corresponding to ∼20 nm spatial resolution, were achieved for both dyes and is comparable to previously reported results.^28^

**Figure 2.**
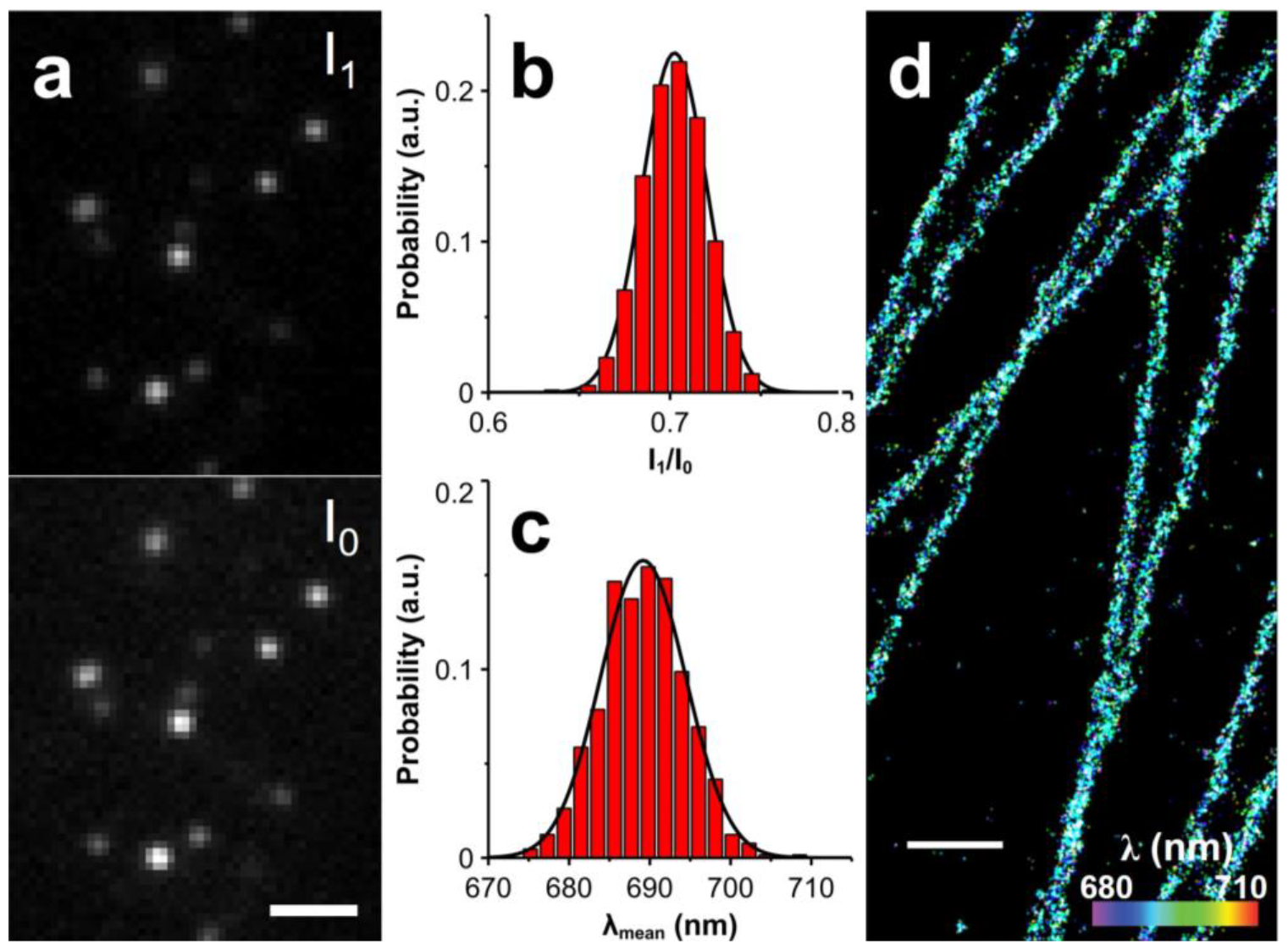
Spectrally resolved single molecule localization microscopy imaging of microtubules in A549 cell. (a) Representative fluorescent images of single Alexa fluor 647 molecules in CGF (I_1_) and reference (I_0_) channels. (b) and (c) show the histogram distribution of I_1_/I_0_ and mean emission wavelength for Alexa fluor 647, respectively. The histogram distribution of mean emission wavelength was fitted by 1D Gaussian function, giving a mean emission wavelength of 689.2 nm and standard deviation (s.d.) of 5.3 nm. (d) Super-resolution fluorescence image of microtubules with color coded spectral mean. Scale bar: 2 μm (a) and 500 nm (d).

### Improved photon budget for SR-SMLM

The achievable spatial resolution in SMLM depends on the collected photons from single fluorophores with a simplified relationship of σ∼ N^-1/2^,^29^ where N is the collected photon number from each fluorophore per emission cycle. For spectral dispersion-based methods (i.e., grating and prism), most of the collected photons are spatially dispersed in the spectral channel to improve the accuracy in wavelength determination. Only a small portion of photons is kept for localizing the positions of molecules, thus limiting the achievable spatial resolution. In our method, the single molecule images in CGF-modified and reference channels are identical except for their brightness. Localized molecules’ positions in both channels can be recombined to improve the spatial resolution with better usage of photon budget. For the case of AF647 dye molecule, ∼ 30% of photon loss was observed in the CGF-modified channel (Figure 2b). By combining photons from both channels for localization, an improvement in spatial resolution by ∼ 1.3-fold (∼ 85% of total photons) can be expected when comparing to that from the reference channel only (∼ 50% of total photons). In comparison to single channel monochromatic SMLM, the spatial resolution is expected to be a ∼ 0.9-fold lower. To test this hypothesis, we imaged AF647 molecules labeled on the microtubules in A549 cell (Figure 3). Conventional fluorescence, single-channel (I_0_), and dual-channel (I_0_ + I_1_) dSTORM images of microtubules in the same regions of cells are presented in Figure 3b-d. We first evaluate the best achievable spatial resolution in dual-channel dSTORM images through cross-section analysis of single microtubule filament (red square in Figure 3c). The normalized histogram distribution of molecule positions is shown in Figure 3e. Fitting the distribution data with 1D Gaussian function gives a full width of half maximum (FWHM) of approximately 62 nm, which is comparable to previously reported results in single color STORM imaging.^28^ Next, we compared the spatial resolution with or without combining photons from CGF-modified (i.e., I_1_) and reference (i.e., I_0_) channels. Collected photons from single AF647 molecules in single and dual channels were measured. As we can see in Figure 3f, the dual-channel provides more photons, which means a higher signal-to-noise ratio can be achieved and images with higher spatial resolution can be obtained. To compare the spatial resolution in single- and dual-channel dSTORM images, we plotted the cross-section profile of molecular positions of two nearby and parallel MT filaments (Figure 3g). Clearly, higher spatial resolution is obtained in the dual-channel dSTORM image as the microtubule filaments are better spatially resolved. The results garnered from these experiments confirm that our methodology can resolve single molecule fluorescence spectra, meanwhile maintaining high spatial resolution through better usage of photon budget.

**Figure 3.**
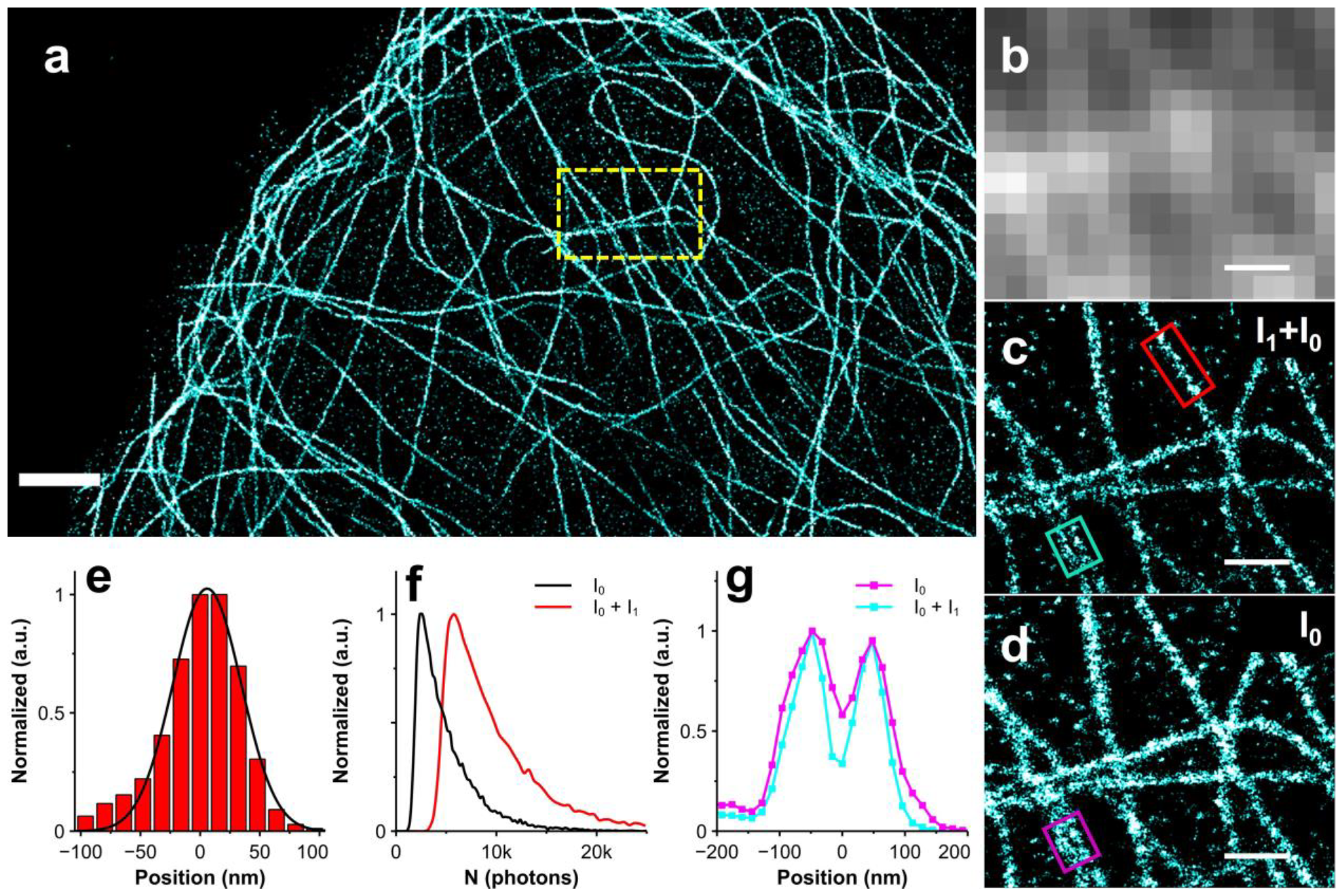
Enhancement of super-resolution imaging resolution by using photons from dual channel imaging. (a) Rendered dSTORM imaging beta-tubulin in A549 cells. Conventional fluorescence image (b), dSTORM image with dual (c) and single (d) channels. (e) Cross-section profile of molecular positions of single MT filament (red box in Figure 3c). The histogram distribution was fitted by 1D Gaussian function, giving a FWHM of 62 nm. (f) Single molecule intensities of Alexa fluor 647 dyes from single channel (black) and dual channel (red). (g) Cross-section profile of molecular positions of two nearby and parallel MT filaments (cyan and magenta box in c and d). Scale bar: 2 μm (a), 500nm (b-d).

### Spectrally resolved super-resolution imaging of subcellular structures

To demonstrate the capability of our method for spectrally resolving dye molecules that overlap heavily in their emission spectra. We double labeled microtubules and mitochondria with AF647 and Cy5.5 respectively. A single 642 nm laser was used to excite and photoswitch all dyes for single molecule imaging. SR-SMLM image of both subcellular features is presented in Figure 4. The color denotes the measured spectral mean of each single dye molecule detected and localized, thus representing their ‘true color’ rather than ‘false color’ assigned. As shown in Figure 4b, different dyes are already readily distinguishable based on their spectral mean distributions without the prior knowledge of spectra of the dyes. Distinct colors are observed for microtubule (cyan) and mitochondria (green) structures. We further quantified the profile of spectral mean at spectrally resolved subcellular features (highlighted in red box in Figure 4b). The histogram distributions of spectral mean for AF647 (microtubule) and Cy5.5 (mitochondria) are shown in Figure 4c. Clearly, they are well resolved from each other. Fitting the distribution data with 1D Gaussian function gives spectral mean of 689.7 ± 7.0 and 707.4 ± 7.5 nm for microtubule (i.e., AF647) and mitochondria (i.e., Cy5.5), respectively. These results indicate that we achieve ultrahigh-throughput and ‘true’ color, spectrally resolved super-resolution imaging of subcellular structures with high spatial and spectral resolution in a simple optical system.

**Figure 4.**
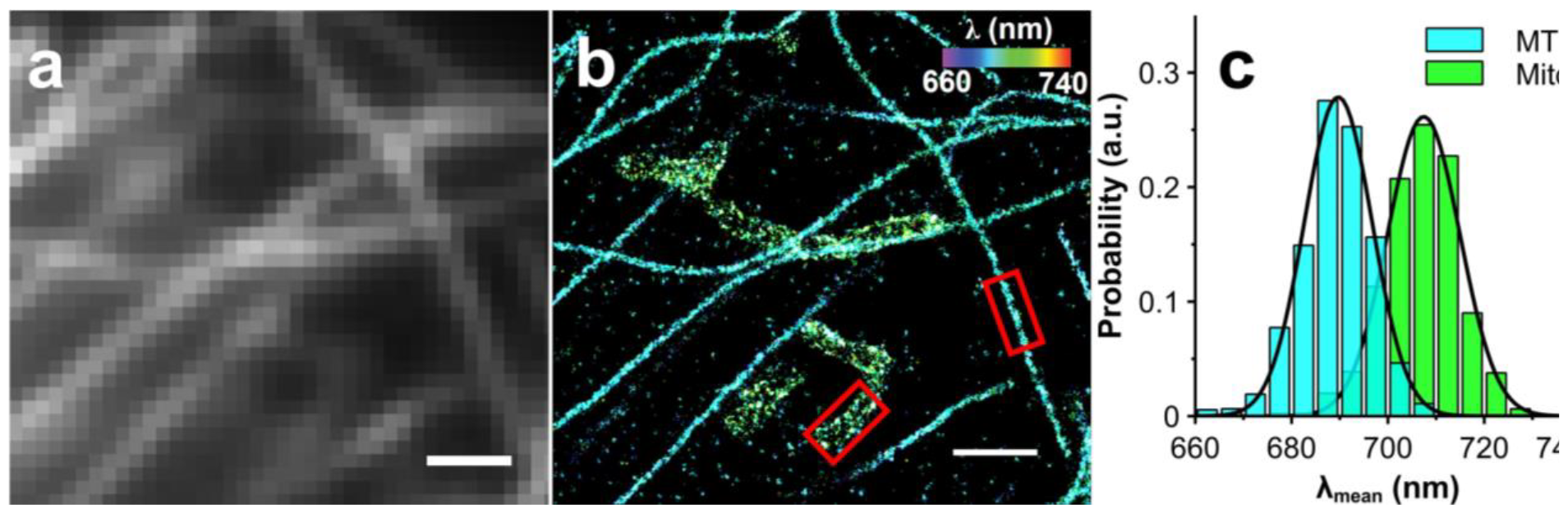
Spectral-resolved super-resolution imaging of subcellular structures. Conventional fluorescence image (a) and spectrally resolved dSTORM image (b) of microtubule and mitochondria. (c) The histogram distribution of measured mean spectral wavelength of AF647 labeled MT and Cy5.5 labeled mitochondria. The distribution data were fitted by 1D Gaussian function, giving the mean spectral wavelength of 689.7 ± 7.0 nm (AF647) and 707.4 ± 7.5 nm (Cy5.5) correspondingly. Scale bar: 1 μm.

## Conclusions

In summary, we present an imaging methodology capable of overcoming challenges associated with current multicolor SMLM techniques. Using a two-channel system employing a simple 50:50 BS and CGF, we can spectrally discriminate between multiple fluorophores and obtain ‘true’ spectral information. By comparing the intensities of single emitters between the modified and unmodified channels, their spectral mean emission is determined at no throughput cost. This methodology enables ultrahigh-throughput imaging, allowing for hundreds of fluorophores in close proximity to be localized with sub-10 nm spatial and sub-5 nm spectral precisions. Because of the simplicity of the setup, this technique can be readily employed in commercial or custom imaging systems with little additional adjustment, making it a generalized and universal technique for unobtrusive multicolor super-resolution imaging. Together, the high-throughput spectral and spatial resolution capabilities paired with the ease of implementation allows this methodology to be a go-to technique for swift SMLM.

## ASSOCIATED CONTENT

### Supporting Information

Experimental details, additional results and discussion, and supporting figures.

## Notes

The authors declare that they have no known competing financial interests or personal relationships that could have appeared to influence the work reported in this paper.

## ACKNOWLEDGEMENTs

This work was supported by the Startup and Honors College Funds from the University of Arkansas Fayetteville.

