## Supplementary Information for "Spectrally Resolved Localization Microscopy with Ultrahigh-Throughput"

### Table of Contents

**S1. Materials and methods** ........ 2

**S2. Supporting figures** ........ 6

### S1. Materials and methods

**Cell culture.** A549 human lung cancer cell line (CCL-185, ATCC) was acquired from the American Type Culture Collection. The A549 cells were cultured in T25 cell culture flask (690160, Greiner Bio-One) in the cell culture medium Dulbecco's Modified Eagle's Medium (DMEM) (10-014-CV, Corning) with 10% fetal bovine serum (FBS) (26140079, Gibco) and 1% Penicillin-Streptomycin (Pen Strep) (15140122, Gibco). µ-slide 8 Well high Glass Bottom chambered coverslip (80807, ibidi) was obtained a gift from ibidi. The chambers were rinsed with PBS before use to remove glass dust. To subculture cells, to each chamber, 50 µL cell suspension solution was added. Then, 250 µL of DMEM containing 10% FBS and 1% Pen Strep was added to every chamber. The chambered coverslip was kept in the cell culture incubator at 37°C with 5% $CO_2$ for 24 hours before the cells were used in imaging experiments.

**Immunostaining of subcellular structures.**

*Materials:*

Cells grown in 8 well-plate.

Pre-warmed Phosphate-buffered saline (BPS)

Pre-warm Fixing Solution (3% v/v Paraformaldehyde (PFA), 0.1% v/v glutaraldehyde (GA), PBS)

Reducing solution (10 mg $NaBH_4$ in 10 mL DI water)

Blocking Buffer (3% w/v Bovine serum albumin (BSA), 0.2% v/v Triton-X100, PBS)

Washing Buffer (0.2% w/v BSA, 0.05% v/v Triton-X100, PBS)

Primary antibody

Dye-conjugated secondary antibody

Storing solution (0.1% v/v Sodium azide, PBS)

*Procedure:*

The Cell culture medium was aspirated from the well-plate chamber and the chamber was washed using 200µL of the warmed PBS. 200 µL of the warmed fixing solution was added to each well-plate chamber. The well plate was left undisturbed for 10 minutes. The fixing solution was aspirated and 200 µL of the reducing solution was added to each chamber. The well-plate was then shaken for 7 minutes using an orbital shaker. For the following steps, the well-plate was shaken for the stated time using an orbital shaker unless stated otherwise. The reducing solution was aspirated and 200 µL of PBS was used to wash the chambers of the well-plate. The washing step was performed 3 times with each washing cycle lasting 7 minutes. The PBS was aspirated and 200 µL of the blocking buffer was added to each chamber of the well-plate. The cells were blocked for 60 minutes. The blocking buffer was aspirated and 150 µL of the primary antibody were added to each chamber of the well-plate. The primary antibody was allowed to incubate for 60 minutes. The primary antibody was aspirated and 200 µL of the washing buffer were added to the chambers of the well-plate. The washing step was performed 3 times with each washing cycle lasting 15 minutes. For the following steps, the well-plate was shielded from light by covering it with aluminum foil. The washing buffer was aspirated and 150 µL of the labeled secondary antibody were added to each chamber of the well-plate. The secondary antibody was allowed to incubate for 45 minutes. The secondary antibody was aspirated and 200 µL of the washing buffer was added to each chamber of the well-plate. The washing step was performed 3 times with each washing cycle lasting 10 minutes. The washing buffer was aspirated and 500 µL of PBS was added to the chambers. This washing step was performed once for 5 minutes. The PBS was aspirated and 200 µL of the fixing solution was added to each well-plate chamber. The well plate was left undisturbed for 10 minutes. The fixing solution was aspirated and 500 µL of PBS was added to each chamber of the well-plate. The washing step was performed 3 times with each washing cycle lasting 10 minutes. The PBS was aspirated and 500 µL of the storage solution was added to each well-plate chamber. The well plate was covered with aluminum foil, and it was stored at 4 $^{o}$C before the imaging experiments.

**Microscope setup.**

The imaging experiments were carried out using a total internal reflection fluorescence microscopy (TIRFM). An adjustable 100-mW 642-nm CW laser (Oxxius, Lannion, France) was focused at the back focal plane of an oil immersion objective (Olympus, UPLAPO100X, NA 1.50) and project a collimated illumination beam into the sample. By shifting the laser beam laterally before entering the objective, the laser beam projects to the sample/coverslip interface at an angle larger than the critical angle. The laser beam was totally reflected, and an evanescent field of the same wavelength was created at the interface. The intensity in the evanescent field decays exponentially as the distance increases from the interface, which effectively excites fluorophores near the interface and increases signal to noise for imaging. A field stop was used to control the illumination area. The fluorescence signal was collected by the same oil immersion objective. A quadra-band filter set (Chroma, TRF89901v2) was used to direct laser for excitation and reject laser background for fluorescence emission. An additional 706/95 bandpass filter (Chroma) was used to further reduce laser scattering background. The collected fluorescence signal was directed into a custom-built dual channel imaging box. The imaging box consists of a 50:50 nonpolarizing beam splitter (Thorlabs), which creates the two channels. In one channel, the fluorescence emission was further modified by a color glass filter (Thorlabs). The modified and nonmodified (i.e., reference) signals were focused on different portions of a highly sensitive EMCCD camera (Andor iXonEM$^+$ Ultra 897 camera) for dual channel imaging.

**dSTORM imaging.**

*Materials:*

GLOX solution (1 mg glucose oxidase, 15.3 µL PBS, 4.7µL of 17 mg/mL catalase)

1M beta-Mercaptoethylamine (MEA)

Imaging Buffer (50mM Tris buffer, 10mM NaCl, 10 % glucose (m/v))

STORM buffer using beta-mercaptoethanol (BME) (1% GLOX (v/v) 1% BME (v/v) in imaging buffer)

STORM buffer using MEA (1% GLOX (v/v) 1% MEA (v/v) in imaging buffer)

*Procedure:*

The storm buffer using BME/MEA was added into the chamber using a micropipette. To exchange the imaging buffers, the original buffer was washed away using PBS buffer. The new imaging buffer was then added into the chamber. Conventional fluorescence image was first taken at low laser power density. Then the laser power was increased to photoswitch the majority of the dye molecules into nonfluorescent dark state. A random subset and spatially resolved dye molecules were kept in fluorescing (on state) in each imaging frame. For each dSTORM imaging experiment, data of 30k-40k frames were collected at an imaging rate of 50 frames per second.

**Data analysis.** The collected single molecule imaging data were analyzed by either ThunderStorm ImageJ plugin or Insight3. The single molecules in both channels were first identified and localized independently. To identify the same single molecule in both channels, a correlation analysis procedure was followed which is shown in Figure S2. We first imaged fluorescent beads on coverslip and use their localized positions to build a transformation matrix. The transformation matrix can then be used to project the spatial coordinates of localized single molecule positions from CGF-modified channel to the refence channel. The projected molecular positions are then compared to those obtained in the refence channel. Molecular positions that are within the same imaging frame and within the localization precision are considered as from the same molecule. Their photon intensities were used for calculating the $I_1/I_0$ ratio. Applying the calibration curve, we obtained the spectral mean ($\lambda_{mean}$). The average localized position of the same molecule in both channels was calculated. Combining the obtained spectral mean and average localized position, spectrally resolved super-resolution image was rendered using Insight3 software. The correlation analysis was done using MATLAB.

### S2. Supporting figures

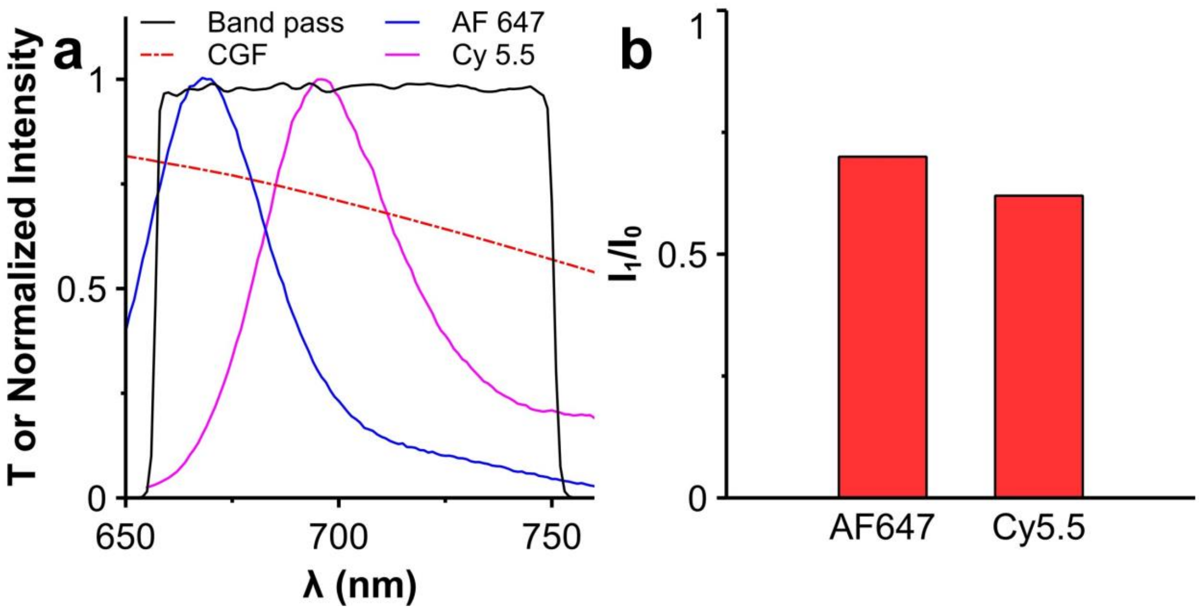


**Figure S1. Fiter set in the dual channel optical imaging system.** (a) Transmission or fluorescence emission profiles of 706/95 band pass, color glass filter, AF647 and Cy5.5. Note that the glass filter only presents in one of the channels. (b) Theoretical intensity ratios in two channels for AF647 and Cy5.5.

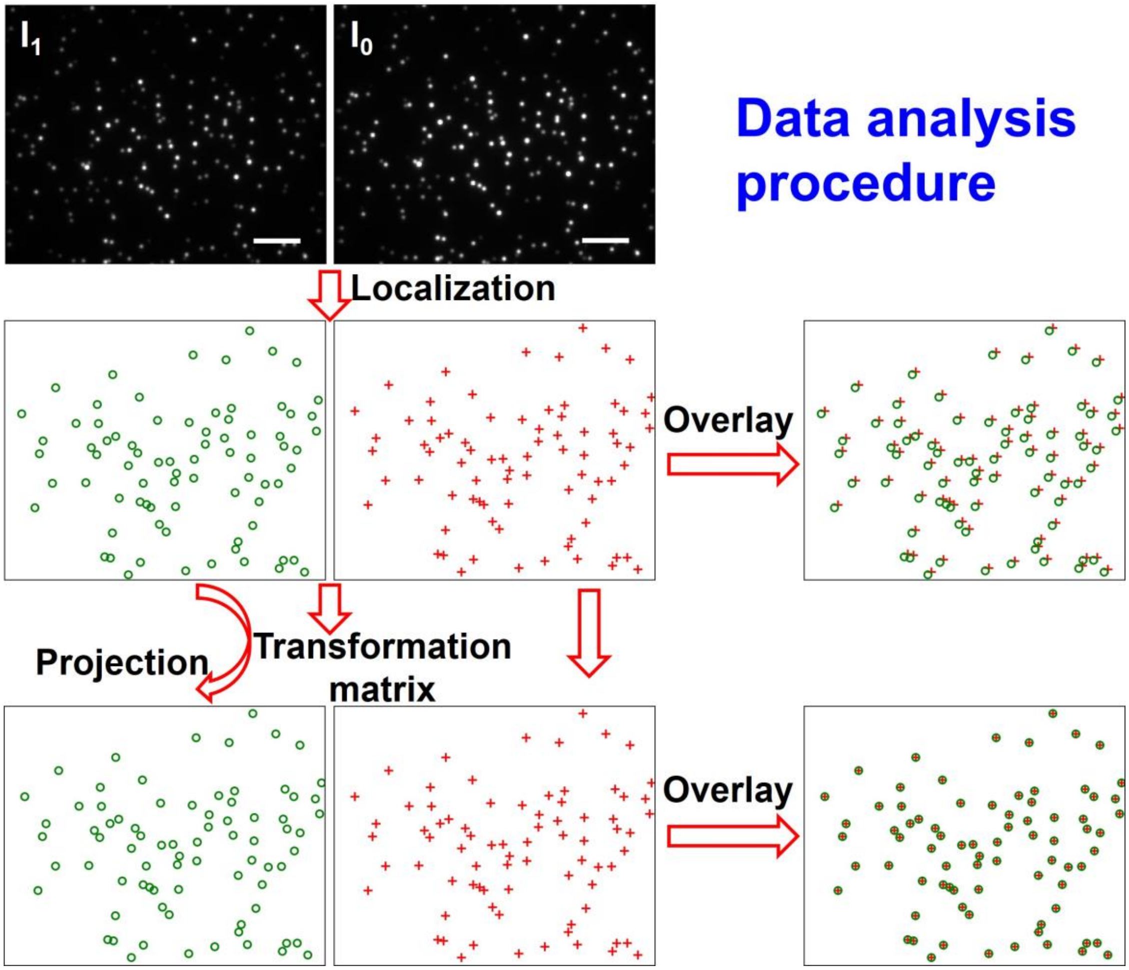


**Figure S2. Schematic view of data processing for identify the same molecule in two channels**. The transformation matrix for the dual channel imaging was calibrated using fluorescent beads first and then applied for correlation analysis of single molecule data.

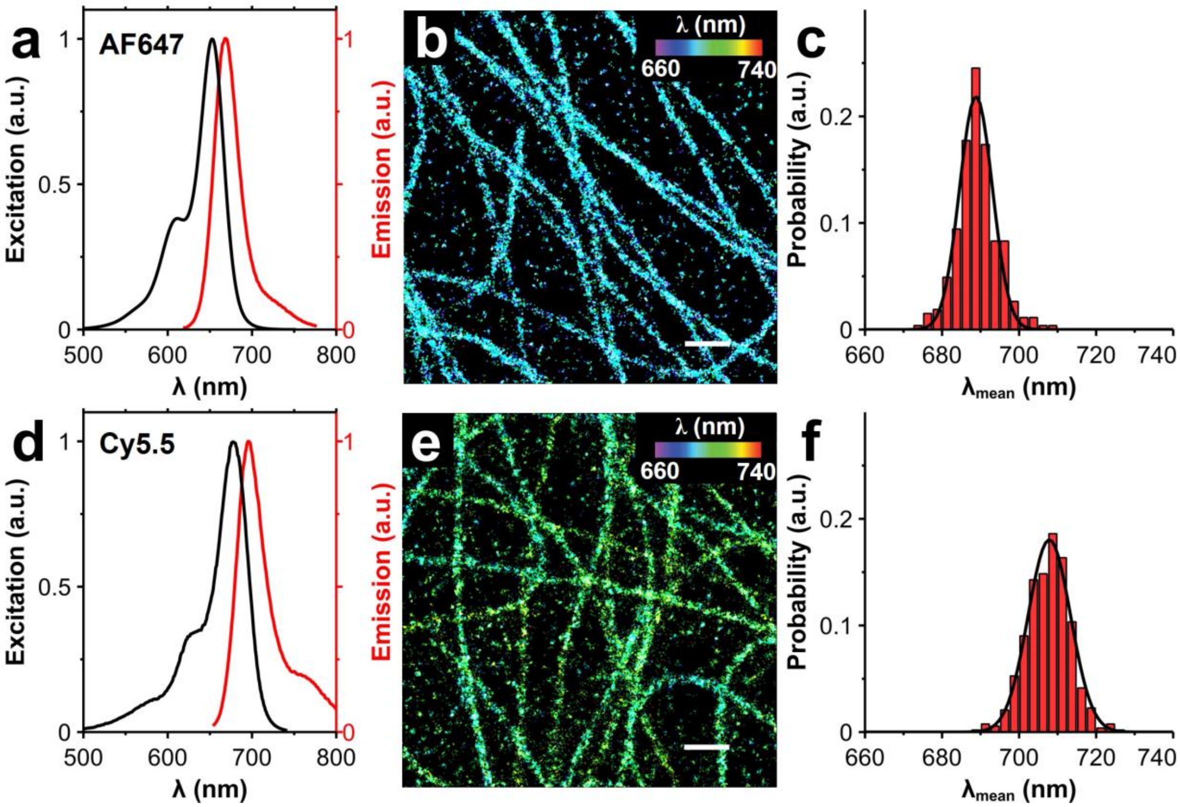


**Figure S3. Spectrally resolved single molecule localization microscopy imaging of microtubules in A549 cell.** (a, d) Excitation and emission spectra of AF647 and Cy5.5. (b, e) Spectrally resolved super-resolution fluorescence image of microtubules labeled with AF647 and Cy5.5. (c, f) The histogram distribution spectral mean for AF647 and Cy5.5. The histogram distribution of spectral mean was fitted by 1D Gaussian function, giving a mean emission wavelength of 689.2 ± 5.3 and 707.9 ± 5.5 nm for AF647 and Cy5.5 respectively. Scale bar: 500 nm.

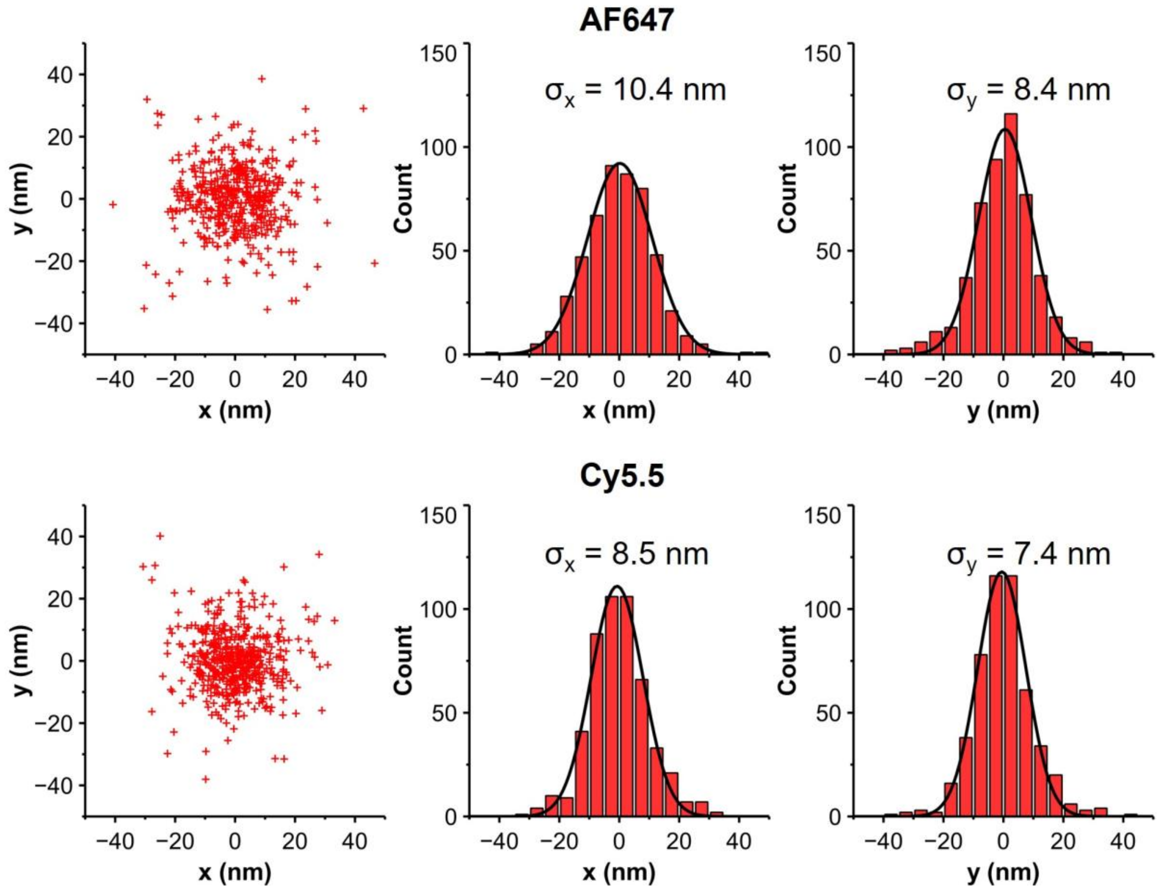


**Figure S4. Localization precisions.** Cluster analysis for quantification of localization precisions using AF647 and Cy5.5.
